# Attentional state, not trait, predicts test performance in video-based learning

**DOI:** 10.1101/2025.05.02.651980

**Authors:** Jens Madsen, Lucas C. Parra

## Abstract

Poor academic performance is often linked to attention deficits, but it remains unclear whether these reflect enduring traits or momentary lapses in focus. In this study, we examined how state and trait-level attention relate to learning from short educational videos. Across four experiments (N = 152), participants completed standardized assessments of inattention, hyperactivity, working memory capacity, and GPA. They then viewed 3-6-minute educational videos while electroencephalography (EEG) tracked neural responses, followed by quizzes assessing short-term retention. Neural synchrony during video viewing, a measure of attentional state, strongly predicted test performance (p < 0.001). In contrast, trait inattention did not predict retention (p > 0.05), although it was negatively associated with GPA. Attentional state was positively associated with working memory capacity but not with trait inattention. These findings suggest that students with attentional difficulties can still learn effectively from short, engaging content, emphasizing the importance of instructional format in supporting diverse learners.

## Introduction

Academic success is shaped by a dynamic interaction of cognitive capacities ^1,2^, individual personality traits ^3^, and educational environments ^4^. Core cognitive functions such as working memory and cognitive flexibility are strong predictors of achievement ^2,5,6^. Among these, attention plays a central role: students who stay focused are better able to process and retain information ^7,8^, whereas attentional lapses disrupt encoding and impair learning ^9^. This is particularly evident in the challenges faced by students with symptoms of attention-deficit/hyperactivity disorder (ADHD), a condition known to affect approximately 5-10% of children and adolescents worldwide ^10,11^. In recent years, ADHD diagnoses have risen globally, a trend attributed to broader diagnostic criteria and increased awareness among clinicians and the public ^12^. ADHD is typically characterized by inattention and hyperactivity-impulsivity, both of which are linked to poor academic outcomes, with inattention showing the strongest associations ^13,14^.

Extensive research has shown that individuals with ADHD, particularly when untreated, tend to perform worse on academic measures such as standardized test scores, grades, and overall school performance, even when these analyses statistically adjust for general cognitive ability (IQ) ^15–17^. This suggests that ADHD impacts learning through mechanisms beyond general cognitive ability, potentially involving chronic attentional disengagement, executive dysfunction, and persistent academic difficulties. Even in undiagnosed individuals, inattention and hyperactivity scores predict school outcomes ^18,19^. These findings align with growing evidence that ADHD traits are best conceptualized as dimensional rather than categorical, reflecting a continuous distribution across the population ^20,21^. This dimensional perspective allows exploration of how these attentional traits impact learning in typical student samples. In line with this dimensional framework, we treat inattention and hyperactivity as continuous, separable trait measures, rather than diagnostic categories, when assessing participants in our general sample.

One cognitive mechanism often implicated in these difficulties is working memory capacity (WMC), a stable trait reflecting how much information an individual can actively maintain and manipulate over short periods of time ^22^. WMC has been shown to independently predict academic achievement across multiple domains, including mathematics ^23^ and reading ^24^. Deficits in WMC contribute to difficulties with multi-step problems, especially when attentional control is required ^23^, and are more pronounced in those with inattention symptoms ^25^, persisting even when IQ and reading ability are controlled ^26^. WMC is a stronger predictor of academic success than IQ in some longitudinal studies ^1^, and underpins higher-order functions such as reasoning, language comprehension, and problem solving ^27^. WMC impairments are associated with inattentive traits, and co-occur with deficits in processing speed and verbal memory ^28,29^.

While attention and memory have traditionally been studied within formal educational contexts, focusing on structured instruction and classroom learning ^30^, students now consume media in radically different formats. Contemporary learners often engage with short-form digital content via platforms like YouTube, TikTok, and social media ^31,32^. This cultural shift raises the possibility that attentional difficulties observed in academic settings may not reflect a general inability to focus, but rather a mismatch between students’ habitual media experiences and the demands of conventional educational formats ^33^. Moreover, frequent media multitasking, encouraged by modern digital platforms, has been associated with fragmented attention and decreased academic performance ^33,34^. This raises an important question: are students able to sustain attention during brief, media-aligned formats?

Short-form educational videos (typically 3-6 minutes) offer a natural test case for this question ^35,36^. These formats may better align with students’ media habits and capture their attention more effectively than traditional lectures. However, some scholars have raised concerns that brief videos promote shallow processing and limit opportunities for deeper learning ^37,38^. One possibility is that attentional difficulties associated with poor academic outcomes may not reflect a general inability to focus, but rather a mismatch between students’ cognitive traits and traditional educational formats. If short videos can support acute attentional engagement, even in individuals with high inattention or hyperactivity, this may suggest they are well suited to diverse learners.

Longitudinal studies link inattention to academic underperformance and self-regulation difficulties ^15,39^, suggesting that inattention traits primarily impairs long-term learning processes. However, recent work documents momentary attentional lapses in brief, structured tasks ^40,41^, raising the possibility that inattention or hyperactivity may also influence learning outcomes even in short-form contexts. Importantly, such traits are distributed continuously across the general population, not limited to clinical diagnoses. This raises an important question: are learning outcomes in short-form video-based instruction more strongly influenced by stable attentional traits (e.g., inattention, hyperactivity, working memory capacity), or by moment-to-moment fluctuations in attentional engagement during the video itself?

To address these questions, we used electroencephalography (EEG) to obtain continuous, objective measures of attentional engagement. Specifically, we analyzed intersubject correlation (ISC) of EEG signals, which quantifies neural response similarity across viewers of the same stimulus ^42,43^. ISC has been shown to increase with attentional focus and to predict individual memory performance following video-based learning ^44,45^.

Our first goal was to characterize how acute distraction alters the strength of neural responses, using a dual-task manipulation in which participants rewatched videos while performing mental arithmetic. Our second goal was to examine how trait-level variables, consistent of inattentive and hyperactive traits as well as, working memory capacity (WMC) relate to attentional engagement, learning outcomes and GPA. Based on prior work, we predicted that fronto-central EEG components would be most sensitive to attentional state and predictive of memory performance ^44,45^. We tested whether inattentive and hyperactive traits would reduce attention and performance, or whether short-form videos remain robust to such variability ^16,17^. We addressed these aims using secondary data analysis of the *BBBD* dataset ^46^, comprising four experiments (N = 152) in which participants viewed short educational videos while EEG was recorded, followed by memory quizzes (administered after each video or block of videos to assess short-term retention), inattention, hyperactivity and WMC assessments, and self-reported GPA. This is the first study to analyze the EEG data from the *BBBD* dataset, and the first to examine neural and trait-level predictors of learning performance using these recordings.

## Results

### What effects does attention have on neural activity?

Prior work has shown that when individuals pay attention during video viewing, their EEG responses become more temporally aligned with those of other viewers. This neural similarity, measured using inter-subject correlation (ISC), is computed by averaging the Pearson correlations between each participant’s EEG time course and those of all other participants. ISC increases with attentional engagement and reliably predicts memory for video content ^44,45^. We first sought to determine which component of the electroencephalography (EEG) signal best reflects attentional modulation, that is, which neural responses show the greatest difference in intersubject correlation (ISC) between attentive and distracted viewing conditions. To do this we analyzed two experiments from the *Brain Body Behavior Dataset* (*BBBD*), comprising 60 participants (age range: 18-57 years; mean age = 25.95; 42 female; see Fig. 1). Participants watched typical short-form informative videos explaining particular topics related to biology or physics. (3-6 minute long) from YouTube. These videos were characterized by their brevity, fast-paced delivery, and dynamic audiovisual style, features common in modern digital platforms such as YouTube and TikTok, and increasingly used in educational settings. After watching the videos in the attending conditions, participants answered quiz questions testing for recall and comprehension. To test the effect of attention on the neural responses, after students had watched the video in the attending condition, they watched the videos again in a distracted condition where they were distracted by carrying out a mental arithmetic task. The mental arithmetic task (counting backward in decrements of 7 from a randomly chosen prime number between 800 and 100) was selected to engage participants’ cognitive resources and divert their attention from the video. This procedure is often used in cognitive neuroscience studies^43^ to ensure that overt behavior of participants remained unchanged and the visual and auditory stimuli remained identical across conditions, thus isolating the effect of attentional engagement.

**Figure 1.**
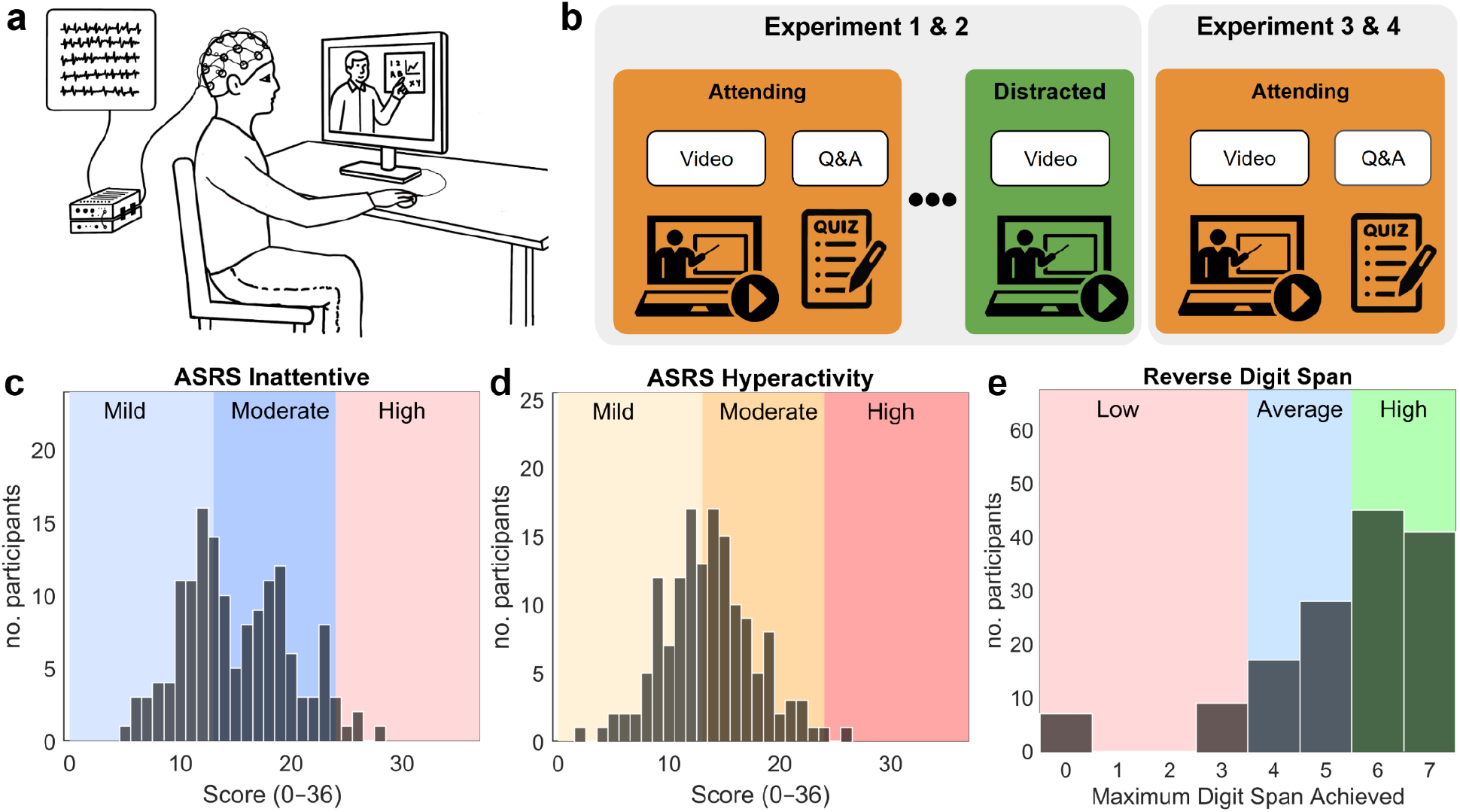
Experimental design and participant characteristics. **a)** Overview of the EEG recording setup during video-based learning tasks. **b)** In Experiments 1 (N=31) and 2 (N=29), participants watched videos under two conditions: attending, where they viewed the videos as they normally would, followed by a quiz; and distracted, where they rewatched the videos while performing a mental arithmetic task. In Experiments 3 (N=43) and 4 (N=49), participants completed only the attending condition, which was originally part of a broader intervention study. Only the non-intervention segments are included in the present analyses. While the procedure was matched across all four experiments, each experiment used a different participant cohort and a partially overlapping set of videos, allowing for assessment of generalizability across stimuli and trait distributions. Additional details, including video overlap and experimental differences, are provided in the Methods section and the *BBBD* dataset paper. **(c, d)** Distributions of inattention and hyperactivity scores on the Adult ADHD Self-Report Scale (ASRS; N=149). Shaded regions in panels (c) and (d) indicate score ranges suggestive of clinically significant symptom levels, as per established guidelines ^47^. **e)** Distribution of maximum reverse digit span scores (N=147), grouped into low (0-3), average (4-5), and high (6-7) working memory capacity ^48^.

We use correlated component analysis (CorrCa ^49^) to extract components that are maximally correlated between subjects. We sort them in order of ISC with the first explaining the strongest correlation between subjects. The first component had a fronto-central distribution on the scalp (Fig. 2), replicating previous results ^44,50,51^. To measure the effect of attention on each component, we compute Cohen’s d-prime for the difference in ISC values between the normal viewing conditions and the subsequent distracted condition (Fig. 2). We find that the first component with the strongest ISC is also most robustly reduced by the distracting task of counting.

**Figure 2.**
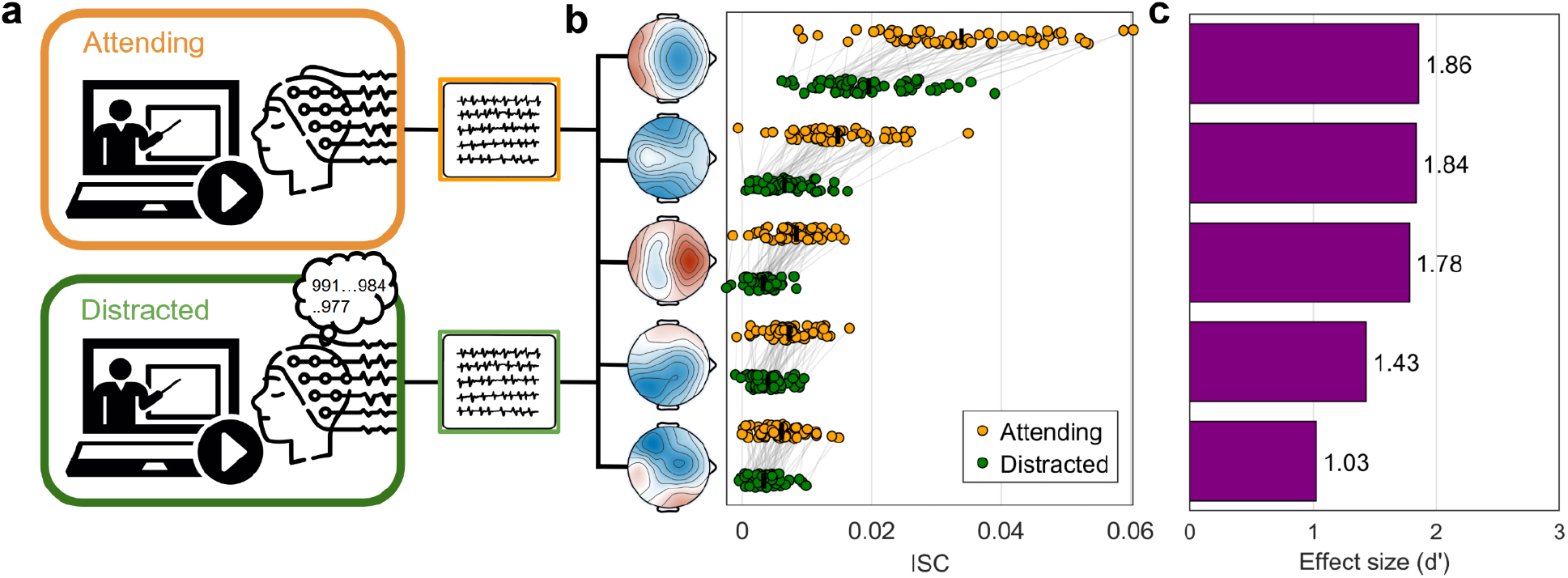
Effect of a distracting secondary task on video-evoked responses (mental arithmetic). **a)** Participants in Experiment 1 & 2 (N=60) rewatched the videos while performing a distracting mental arithmetic task (counting backwards silently in their heads from a random prime number between 800-1000 in steps of 7). **b)** EEG data were analyzed using correlated component analysis (CorrCA), resulting in 5 spatial filters (C1-5). We compute the ISC for each of these components in the distracted and attending conditions. Each dot is a participant and lines connect each participant’s ISC values. **c)** Bar plots indicate the effect size (Cohen’s *d′*) on ISC. Positive d’ indicates a drop in ISC in the distracted conditions. All differences were statistically significant (paired t-test, p < 10^−8^, N=60, d-prime here is the t·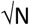^52^).

### Does attention or traits affect students’ test scores?

Next, we analyzed the factors that predict quiz performance. The quizzes consisted of multiple choice questionnaires (10-12 questions per video) for each of the different 11 videos that were used across the four experiments from *BBBD*, comprising 152 participants (age range: 18-57 years; mean age = 25.58; 93 female). These quizzes were completed immediately after each video (or block of videos), providing a measure of short-term retention. To capture student traits, prior to the video watching, we asked them to complete a Working Memory Capacity test (reverse Digit span test). In this test the participants are shown a sequence of numbers (one-by-one) and are then asked to write the numbers in reverse order. In addition, participants filled out the Adult ADHD Self-Report Scale ^53^ which gives a score for *inattentiveness* and *hyperactivity* (see scores in Fig. 1), and we also asked them to report their GPA, as a measure of overall academic performance. We used a generalized linear model to determine how attention, measured as ISC, as well as working memory capacity, GPA, inattentiveness and hyperactivity affect performance. We find that GPA and WMC are predictors of test scores (*t*(109)=3.41, *p*=9·10^−4^, *t*(109)=3.374, p=0.001, see Supplement S1 for complete table of statistical analysis), but inattention and hyperactivity are not (*p*>0.05) (Fig. 3a). Attention, as measured with the ISC of the first fronto-central component is a strong predictor of test score (*t*(109)=4.11, p=7.67·10^−5^).

**Figure 3.**
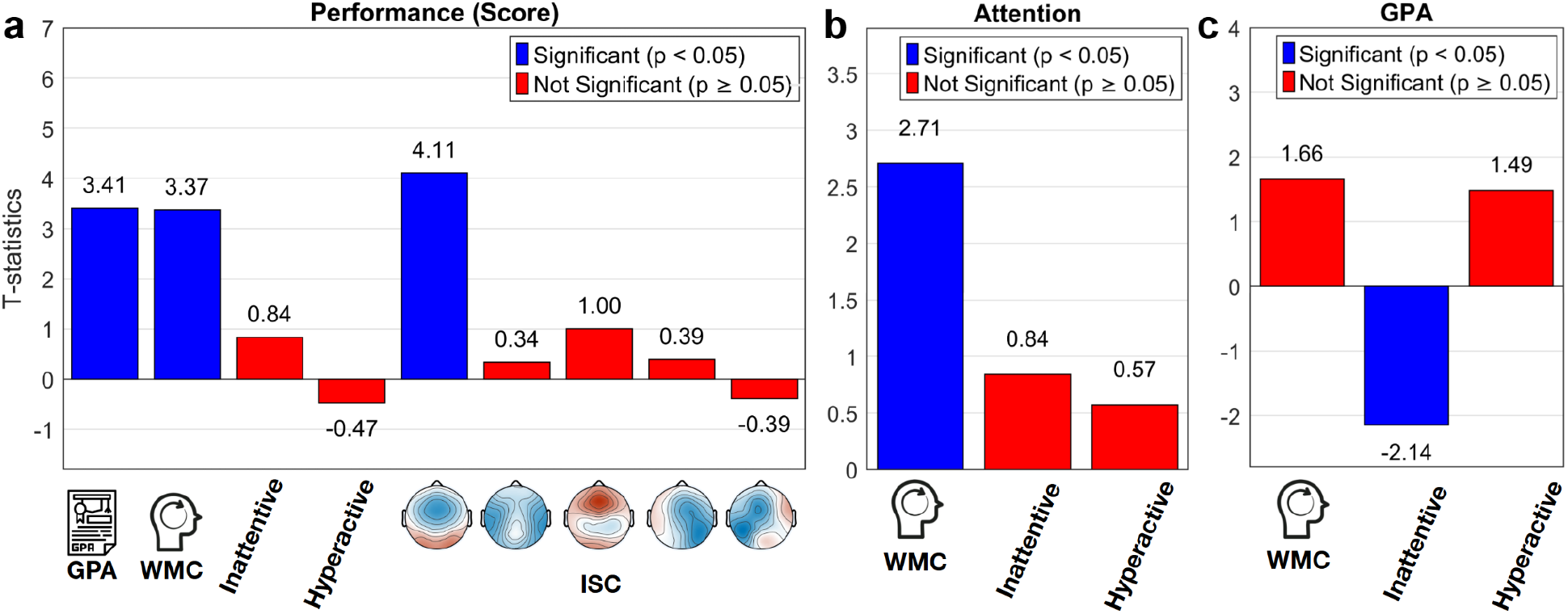
Factors that affect learning performance. **a)** t-statistics for the generalized linear model, modeling the behavioral performance of students answering questions about each short-form educational video (10-12 questions per video, N=152 participants). **b)** t-statistics for the GLM modeling the first ISC-EEG component. **c)** t-statistics for the GLM modeling GPA.

### Mediation analysis of the interaction between performance, attention and traits

The previous analysis considered long-term traits, and the current state of the student, i.e. acute attentional state while watching video (ISC) and current overall academic performance (GPA). It is conceivable that the trait factors affect the current state, and thus, that they affect performance indirectly via GPA or ISC. To perform this type of mediation analysis ^54^ we assume, a-priori, that long-term traits affect the current state, but not vise-versa (we will discuss the basis for this and alternatives assumptions in the Discussion section). We therefore compute a model for ISC and GPA with traits as predictors (WMC, Inattention, Hyperactivity), as summarized in Fig. 4. For attention we focus on ISC of the first fronto-central component as it was most strongly modulated by attention, and was the main predictor of performance. ISC of this component was predicted by WMC with a positive effect (*t*(142)=2.707, *p*=0.0076, see Fig. 3b), but not by inattention or hyperactivity traits (*p*>0.05, the full statistical table is in the Supplement S1). GPA was predicted by inattention with a negative effect (*t* (115)=-2.14, *p*=0.034, see Fig. 3c) but not by WMC or hyperactivity (p>0.05, the full statistical table is in the Supplement S1).

**Figure 4.**
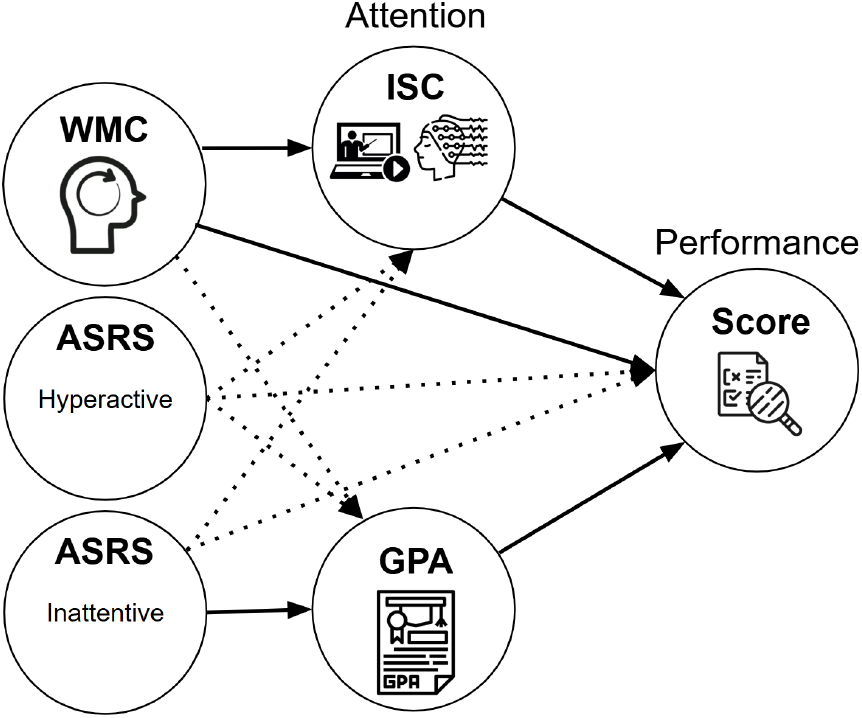
Summary of results: In this mediation analysis, all possible effects from left to right were tested. Solid arrows indicate statistically significant effects, dashed arrows means there was no significant effect.

## Discussion

The present study investigated how acute attentional engagement and individual trait measures predict learning outcomes from short-form educational videos. Our findings demonstrate that the strongest predictor of immediate memory performance was acute attention to the video, as indexed by intersubject correlation (ISC) of EEG signals. This result aligns with foundational theories that posit attention as a *sine qua non* for effective encoding and learning ^55,56^. Moreover, our neural measure of attention has previously been shown to predict quiz performance even weeks after exposure to educational material ^44,57^, underscoring the robustness of ISC as a marker of attentional engagement with dynamic stimuli.

Beyond acute attention, long-term measures such as working memory capacity (WMC) and GPA emerged as significant predictors of performance. This finding is consistent with extensive literature demonstrating that WMC underpins academic achievement across diverse domains, including mathematics and reading comprehension ^2,23,24^. WMC facilitates the maintenance and manipulation of information during learning tasks ^22^, and our results suggest it also supports moment-to-moment attentional engagement with educational stimuli. Higher GPA, a proxy for overall academic success, likely reflects accumulated cognitive skills and prior knowledge that aid in encoding new information, consistent with theories of educational attainment ^58^.

In contrast, self-reported inattentiveness, while a predictor of lower GPA, did not significantly predict neural attention or immediate memory performance. This finding supports the view that ADHD traits, particularly inattention, impact cumulative academic outcomes through chronic disengagement ^16,17^, rather than through acute deficits in attentional engagement with short, structured materials. Notably, neither inattentiveness nor hyperactivity significantly impaired attention to or learning from short videos, suggesting that modern short-form educational content can successfully engage students across a range of attentional profiles.

Our EEG results further corroborate prior work demonstrating that attentional modulation during naturalistic narrative engagement (i.e., non-synthetic educational videos drawn from real-world platforms and widely viewed by online audiences) is predominantly reflected in fronto-central neural components ^43,50^. Here we find that it is this EEG component that best predicts learning performance. The spatial EEG component with an anterior-central topography showed the strongest intersubject correlation and was the best predictor of attention and learning. This pattern is consistent with large-scale attentional control networks implicated in cognitive neuroscience ^56,59^, and supports the interpretation that synchronized neural activity in fronto-central regions reflects increased levels of arousal and autonomic function ^50^.

This conclusion is also consistent with the positive effect we found of WMC onto ISC. This analysis was motivated by the fact that WMC reflects a general cognitive skill of an individual. But of course, WMC could be aided by an ability to maintain attention, and an ability to retrain content in working memory may help maintain attention. So these two factors likely interact, and we could have changed the direction of our analysis without affecting any of the other results.

Our study also provides broader insight into contemporary educational environments. As students increasingly consume fast-paced digital content, short-form educational videos appear well-suited to engage attention even among individuals with attentional trait deficits. This suggests that concerns about a “crisis of attention” in younger generations ^31,32^ should be nuanced: when content matches habitual media consumption patterns, acute attentional engagement remains normal. However, we caution that while acute attention predicts immediate memory performance, it does not necessarily guarantee deeper semantic learning or long-term consolidation into broader knowledge networks, as emphasized in the levels-of-processing framework ^60^. Capturing attention is necessary but insufficient for durable learning and comprehension. Future educational strategies should not only harness the engagement potential of short-form media but also incorporate scaffolds that promote elaborative, semantic processing, encouraging integration of new material into existing cognitive frameworks ^61^.

In conclusion, our results highlight the critical role of acute attentional engagement in video-based learning and suggest that trait-level inattentiveness may not necessarily impede learning when materials are brief, structured, and engaging. As education continues to evolve in an increasingly digital landscape, understanding the interaction between neural attention, cognitive traits, and media formats will be essential for designing effective, inclusive educational strategies.

### Limitations of the study

Several limitations should be noted. First, our study focused on immediate retention after video viewing; it remains unknown whether acute attentional engagement predicts long-term knowledge retention or transfer to novel contexts. Second, our findings are based on brief, single-session interventions and may not generalize to more extended or cumulative learning scenarios. Future research should address these questions through longitudinal designs and by exploring interventions that foster deeper processing during video-based learning. Third, we used only short-form educational videos, common in online platforms. It remains unclear whether the observed relationships extend to longer instructional materials, where sustaining attention is more difficult and mind-wandering more likely ^62^.

Fourth, the videos likely varied in their motivational appeal, which may have influenced attention and retention. Although we did not manipulate this directly, future studies could systematically vary emotional or motivational content. For example, self-directed video selection during study may enhance motivation and attention compared to researcher-selected topics ^63^.

Lastly, the analysis of GPA was based on self-reports and only applicable to college students that had a GPA. Indeed, several participants (42 out of 152) did not report GPA which may have been a bias against reporting low scores, or they may not have had a GPA as they did not attend college. So generally, the GPA analysis should be interpreted with caution.

### STAR Methods

#### Resource availability

##### Lead Contact

Jens Madsen

##### Materials availability

This study did not generate new unique reagents or materials.

##### Data and code availability

The code to reproduce each figure is available on osf.io/u72k3. The data used for the experiment is publically available at *BBBD* website bbbd.pythonanywhere.com. The spatial filters used in the analysis are shared on OSF as well.

## Experimental Model and Subjects

### Human subjects

A total of 152 participants from four experiments (Experiments 2-5 of the Brain, Body, Behavior Dataset [BBBD]) were included in this study. All participants watched short, informative audiovisual narratives while EEG was recorded, along with other physiological and behavioral measures depending on the experiment. Participants provided informed consent under protocols approved by the relevant ethics committees. In Experiment 2, 31 participants were included (21 female; age range = 18-50 years, M = 25.97, SD = 8.26). In Experiment 3, 29 participants were included after removing 3 due to poor data quality (21 female; age range = 18-57 years, M = 25.93, SD = 8.94). In Experiment 4, 43 participants were included after removing 3 (24 female; age range = 18-45 years, M = 25.09, SD = 6.49). In Experiment 5, 49 participants were included after removing 2 (27 female; age range = 18-49 years, M = 25.55, SD = 7.97). In total, 152 participants (93 female) aged 18-57 years (M = 25.58, SD = 7.77) were analyzed across the four experiments. Demographic and modality information is detailed in the *BBBD* dataset documentation. Participants were primarily college students or recent graduates recruited from the City College of New York or people from the surrounding community. While we did not collect self-reported race or ethnicity, the City College of New York serves one of the most diverse student bodies in the United States, and our sample reflects this broader population.

### Stimuli

Across the 4 experiments participants watched a subset of 11 unique informative audiovisual narratives selected from popular YouTube channels known for short explanatory content, including minutephysics, Kurzgesagt - In a Nutshell, MinuteEarth, What If, Be Smart, Khan Academy, and eHowEducation. These videos cover topics in physics, biology, astronomy, and computer science and have been used in prior studies investigating attention and learning from video-based instruction. In Experiment 2, participants viewed five videos from minutephysics, Kurzgesagt, and MinuteEarth, covering topics in physics, biology, and computer science. Video durations ranged from 2:23 to 7:49 minutes (M = 4.94 min). In Experiment 3, six videos were shown, sourced from Khan Academy, eHowEducation, What If, and Be Smart. Topics included biology, physics, and astronomy, with durations ranging from 3:12 to 6:27 minutes (M = 5.16 min). In Experiment 4, six videos overlapped with prior experiments, including content from minutephysics, Kurzgesagt, What If, Be Smart, and eHowEducation, covering physics and biology topics (range: 3:12-7:49 minutes). In Experiment 5, participants viewed four videos from minutephysics, Kurzgesagt, and Be Smart, with durations between 3:29 and 7:49 minutes (M = 5.88 min). All videos were audiovisual narratives that presented core scientific or conceptual explanations in a visually engaging style. The full list of videos, their durations, source channels, and YouTube IDs are documented in the *BBBD* dataset paper.

### Procedure

All experiments were conducted at the City College of New York in a sound-attenuated booth with controlled ambient lighting. The study protocol was approved by the Institutional Review Boards (IRB) of the City University of New York, and documented informed consent was obtained from all participants. Participants were seated approximately 60 cm from a 27” monitor, and videos were presented with audio played through stereo speakers positioned at 60° angles on either side of the screen. EEG and EOG signals were recorded throughout the experiments using a BioSemi ActiveTwo system. Additional physiological and behavioral signals (e.g., ECG, gaze, pupil diameter, head position, respiration) were also collected and are available in the full *BBBD* dataset, but are not analyzed in the current study.

In all experiments, participants watched informative audiovisual narratives under an attentive condition, in which they were instructed to view the videos as they normally would at home while remaining relaxed and still. After each video (or set of videos), participants completed a multiple-choice quiz (10-12 items per video) testing for recall and comprehension of the content. The order of videos and questions was randomized across participants to control for order effects.

Experiments 3 and 4 (i.e., Experiments 4 and 5 in the BBBD dataset) were originally part of a separate intervention study. Only the non-intervention segments were analyzed here, using the same attending procedure as in the other experiments. These experiments were included to expand trait variability and stimulus diversity. While some videos overlapped across experiments, others differed, supporting generalizability across participants and content.

In Experiments 1 and 2 (corresponding to *BBBD* Experiments 2, 3), participants also viewed the same videos under a distracted condition, where they were asked to silently count backwards in decrements of 7 from a randomly selected prime number between 800 and 1,000. This condition was designed to suppress attention using a mental arithmetic task.

The segmentation of physiological recordings was aligned precisely across participants using embedded audiovisual markers (a flash and beep) placed at the beginning and end of each video, detected via a StimTracker (Cedrus).

### Inattention and Hyperactivity

To assess trait inattention and hyperactivity participants completed the 18-item Adult ADHD Self-Report Scale (ASRS^53^), a widely used self-report instrument assessing symptoms of inattention and hyperactivity in adults. The ASRS consists of two 9-item subscales: one measuring inattentive symptoms and the other hyperactive/impulsive symptoms. Each item was rated on a 5-point Likert scale (1 = never, 5 = very often). Subscale scores were computed by summing the responses for the respective items, yielding a total score for inattention and for hyperactivity (see the distribution of ASRS subscale scores Fig. 1c-d). These sum scores were treated as continuous variables in all statistical models. Shaded regions in the figure reflect score ranges commonly interpreted as indicative of clinically significant symptom levels based on published guidelines ^47^.

### Grade Point Average (GPA)

Grade Point Average (GPA) was self-reported by participants who were current or former college students (N = 123), using a standard 0-4 scale. Participants who had not attended college (N = 29) did not report GPA and were excluded from GPA-related analyses. For 13 participants with missing GPA data, the sample median GPA was imputed. For modeling purposes, GPA scores were normalized to the 0-1 range, and a small number of self-reported values exceeding 4.0 were clipped to the upper bound. GPA served as a proxy for broader academic achievement in subsequent analyses.

### Working Memory Capacity (WMC)

WMC is often assessed with the Reverse Digit Span Task. In this task participants are asked to repeat sequences of digits in reverse order. Sequence lengths ranged from 3 to 7 digits, with two trials per length, yielding a total of 10 trials. Each response was scored as correct if all digits were recalled in the correct reversed order. To quantify performance, we computed a maximum span score, defined as the maximum span length at which the participant correctly recalled at least one of two trials. This scoring approach is consistent with standard procedures used in neuropsychological assessments, where maximum span length reflects working memory capacity ^48^. This maximum span score served as a continuous index of working memory capacity and was used in all statistical models.

## Quantification and Statistical Analysis

### Preprocessing of EEG

The EEG was recorded using a BioSemi Active Two system at a sampling frequency of 2048 Hz. Participants were fitted with a standard, 64-electrode cap following the international 10/10 system. In addition the electrooculogram (EOG) was recorded with six auxiliary electrodes (one located dorsally, ventrally, and laterally to each eye).

The EEG and EOG data was first filtered with an analog band-pass filter between 0.016 and 250 Hz by the Active two system. The signal was then digitally high-pass filtered (0.5 Hz cutoff) and notch filtered at 60 Hz (line noise). To remove artifacts and outliers Robust PCA was used, and subsequently the signal was low-pass filtered (64 Hz cutoff) and down-sampled to 128 Hz. Bad electrode channels were identified manually and replaced with interpolated channels. The interpolation was performed using the 3D cartesian coordinates from the electrode cap projected onto a plane. An interpolation method was used to recreate each channel, derived from all surrounding “good” electrodes. The EOG channels were used to remove eye-movement artifacts by linearly regressing them from the EEG channels, i.e. least-squares noise cancellation.

In each EEG channel, outlier samples were identified (values exceeding 4 times the distance between the 25th and the 75th quartile of the median-centered signal) and samples 40 ms before and after such outliers were replaced with interpolated samples.

### Intersubject correlation and attention analysis of EEG

For the neural signals recorded using EEG the Inter-subject correlation was computed using correlated component analysis ^49^. This method finds a model that linearly combines electrodes that capture evoked activity most correlated between subjects. The model consists of several projection vectors that linearly combine electrodes, on which the data is projected (components), capturing independent sources of correlation between subjects. The ISC of each component is obtained by computing the correlation coefficients of the projected EEG time-course between each participant and all other participants. In order to compare ISC values across the 4 different experiments, spatial filters were estimated on Experiment 3 and applied to all other experiments.

For the analysis of on attentional modulation of ISC, we computed the model combining data from Experiments 1 and 2 including the attending and distracted conditions. For the analysis of quiz performance and mediation, we computed the components with the attentive conditions of experiment 2, and used this model to measure ISC in the attentive conditions for all 4 experiments.

### Statistical analysis

We used generalized linear mixed effect models following the hypothesized causal effects shown in Fig. 4 with all arrows considered in the models pointing from left to right. In abbreviated Wilkson notation used by the matlab function fitglme() the models read as follows:

To model the factors affecting quiz performance, i.e. number of correct responses:

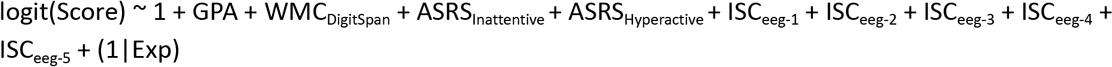

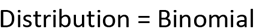

To model factors affecting ISC for the first component of the EEG:

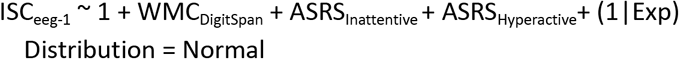

To model the factors affecting GPA:

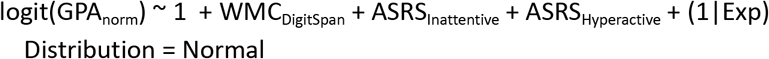

Because quizzes, videos and sample populations differed across experiments we included Exp as a categorical random effect variable in all models. All other factors were fixed effect graded variables. The model for correct responses assumes a Binomial distribution and the fit is given the possible maximum correct responses which differs across quizzes. To allow the use of the logistic link function, GPA was normalized to 0-1, with values exceeding 1 clipped (a few participants reported GPA above 4).

## Author Contributions

J.M. and L.C.P. contributed to the conception and design of the study. J.M. contributed to the acquisition of data. J.M. contributed to the analysis of data. J.M. and L.C.P. contributed to the statistical analysis of the data. J.M. and L.C.P. contributed to the writing of the manuscript and/or the figures.

## Declaration of generative AI and AI-assisted technologies

During the preparation of this work, the author(s) used ChatGPT in order to correct language, grammar and spell check. After using this tool or service, the author(s) reviewed and edited the content as needed and take(s) full responsibility for the content of the publication.

## Supplemental information

### Section S1 - Generalized linear model output

#### Factors affecting performance (score)

Generalized linear mixed-effects model fit by PL

Model information:

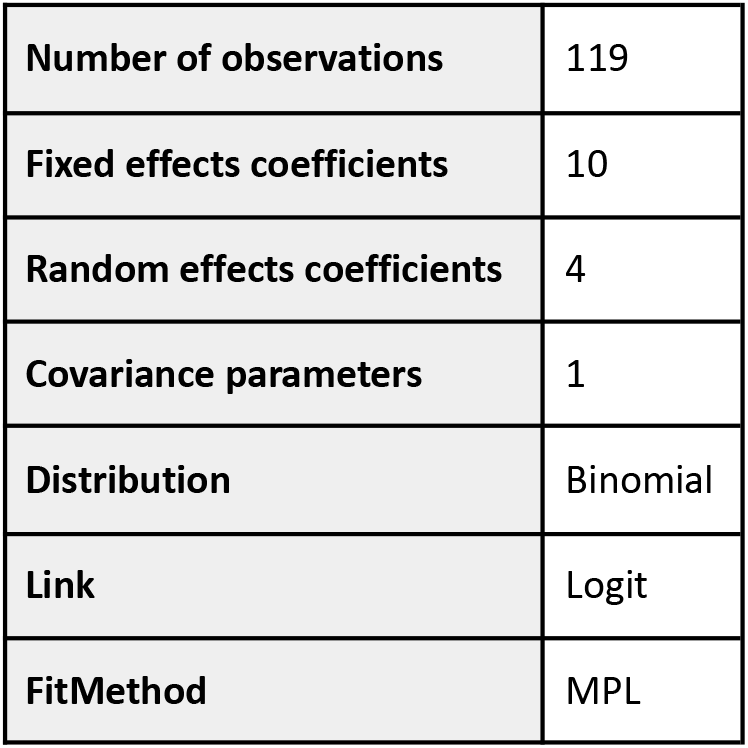

##### Model Formula

**Outcome:**

Score

**Fixed Effects:**

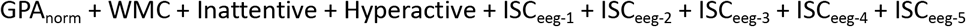

**Random Effects:**

(1 | Exp)

**Full Model Formula:**

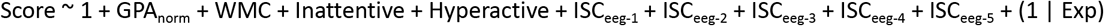

Model fit statistics:

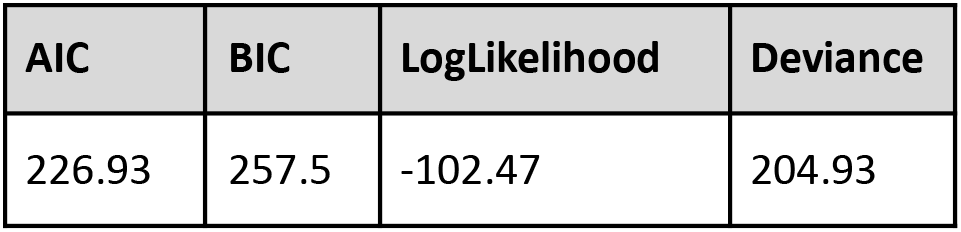

| AIC | BIC | LogLikelihood | Deviance |
| --- | --- | --- | --- |
| 226.93 | 257.5 | -102.47 | 204.93 |

Fixed effects coefficients (95% CIs):

| Predictor | Estimate | SE | t-Stat | DF | p-Value | CI Lower | CI Upper |
| --- | --- | --- | --- | --- | --- | --- | --- |
| (Intercept) | -1.8019 | 0.32212 | -5.5939 | 109 | 1.67E-07 | -2.4404 | -1.1635 |
| GPA | 0.20405 | 0.059833 | 3.4103 | 109 | 0.000911 | 0.085458 | 0.32263 |
| WMC | 0.067786 | 0.02009 | 3.374 | 109 | 0.001027 | 0.027967 | 0.1076 |
| Inattentive | 0.005509 | 0.006586 | 0.83649 | 109 | 0.40471 | -0.00754 | 0.018562 |
| Hyperactive | -0.00369 | 0.007807 | -0.47325 | 109 | 0.63698 | -0.01917 | 0.011779 |
| ISC<sub>eeg-1</sub> | 19.474 | 4.7377 | 4.1105 | 109 | 7.67E-05 | 10.084 | 28.864 |
| ISC<sub>eeg-2</sub> | 2.3485 | 6.9659 | 0.33714 | 109 | 0.73666 | -11.458 | 16.155 |
| ISC<sub>eeg-3</sub> | 11.688 | 11.633 | 1.0047 | 109 | 0.31728 | -11.369 | 34.745 |
| ISC<sub>eeg-4</sub> | 5.1097 | 13.017 | 0.39254 | 109 | 0.69542 | -20.689 | 30.909 |
| ISC<sub>eeg-5</sub> | -4.8603 | 12.542 | -0.38753 | 109 | 0.69912 | -29.718 | 19.997 |

Random effects covariance parameters:

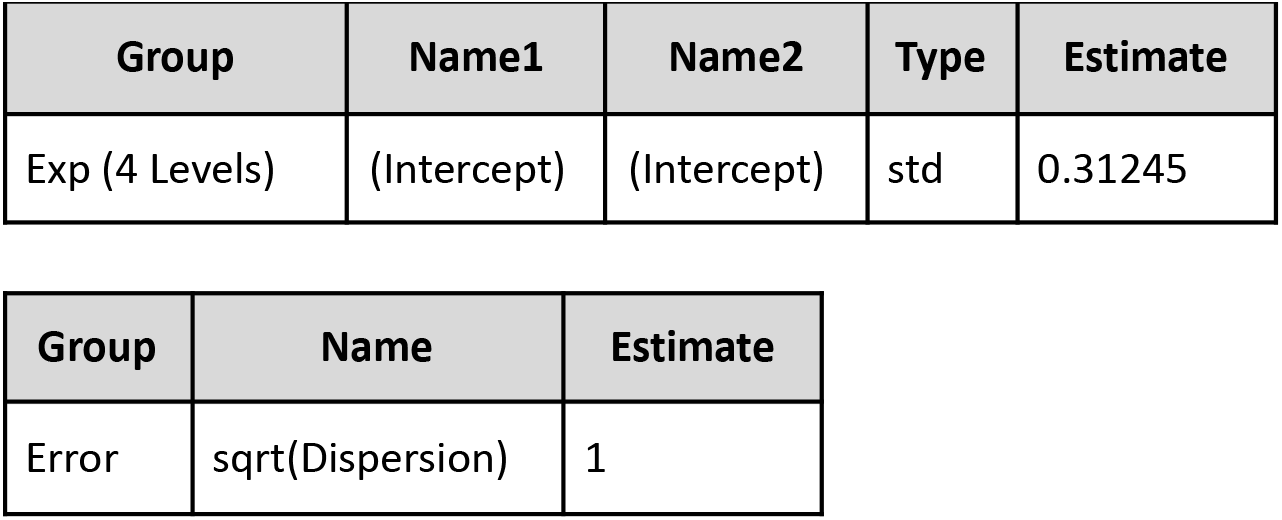

| Group | Name1 | Name2 | Type | Estimate |
| --- | --- | --- | --- | --- |
| Exp (4 Levels) | (Intercept) | (Intercept) | std | 0.31245 |

| Group | Name | Estimate |
| --- | --- | --- |
| Error | sqrt(Dispersion) | 1 |

#### Factors affecting ISC

Generalized linear mixed-effects model fit by PL

Model information:

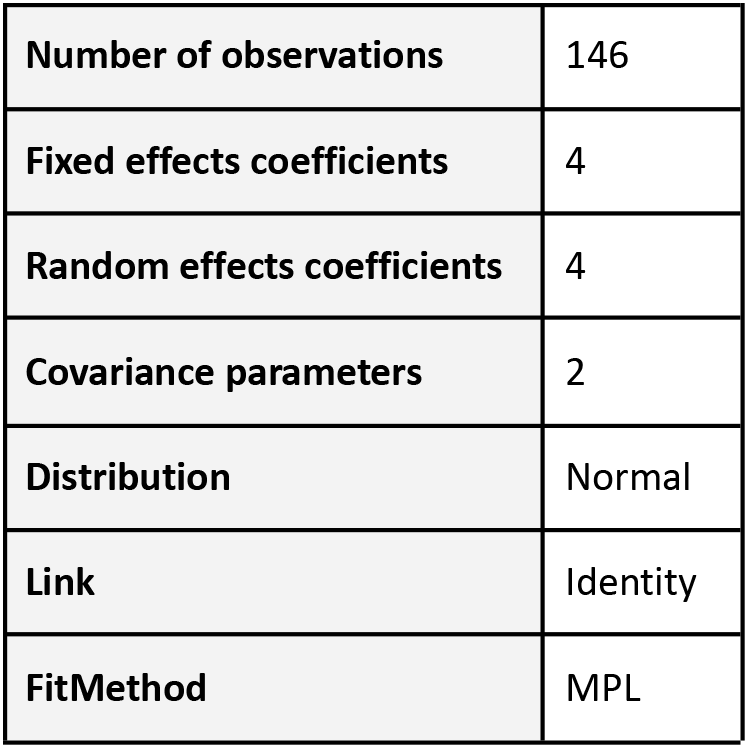

##### Model Formula

**Outcome:**

ISC_eeg-1_

**Fixed Effects:**

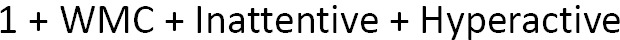

**Random Effects:**

(1 | Exp)

**Full Model Formula:**

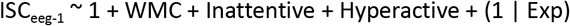

Model fit statistics:

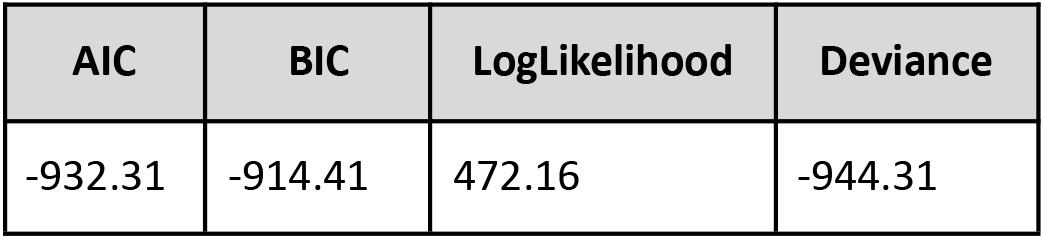

| AIC | BIC | LogLikelihood | Deviance |
| --- | --- | --- | --- |
| -932.31 | -914.41 | 472.16 | -944.31 |

Fixed effects coefficients (95% CIs):

| Predictor | Estimate | SE | t-Stat | DF | p-Value | CI Lower | CI Upper |
| --- | --- | --- | --- | --- | --- | --- | --- |
| (Intercept) | 0.022575 | 0.0046203 | 4.8861 | 142 | 2.74e-06 | 0.013442 | 0.031709 |
| WMC | 0.0012598 | 0.0004655 | 2.7067 | 142 | 0.00763 | 0.0003397 | 0.00218 |
| Inattentive | 0.0001427 | 0.0001695 | 0.8419 | 142 | 0.40129 | -0.0001924 | 0.0004777 |
| Hyperactive | 0.0001129 | 0.0001971 | 0.5726 | 142 | 0.56782 | -0.0002768 | 0.0005025 |

Random effects covariance parameters:

| Group | Name1 | Name2 | Type | Estimate |
| --- | --- | --- | --- | --- |
| Exp (4 Levels) | (Intercept) | (Intercept) | std | 0.0052881 |

| Group | Name | Estimate |
| --- | --- | --- |
| Error | sqrt(Dispersion) | 0.0092081 |

#### Factors affecting GPA

Generalized linear mixed-effects model fit by PL

Model information:

|  |  |
| --- | --- |
| **Number of observations** | 119 |
| **Fixed effects coefficients** | 4 |
| **Random effects coefficients** | 4 |
| **Covariance parameters** | 2 |
| **Distribution** | Normal |
| **Link** | Logit |
| **FitMethod** | MPL |

##### Model Formula

**Outcome:**

GPA_norm_

**Fixed Effects:**

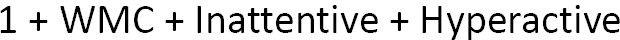

**Random Effects:**

(1 | Exp)

**Full Model Formula:**

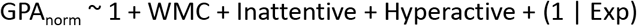

Model fit statistics:

| AIC | BIC | LogLikelihood | Deviance |
| --- | --- | --- | --- |
| 316.2 | 332.87 | -152.1 | 304.2 |

Fixed effects coefficients (95% CIs):

| Predictor | Estimate | SE | t-Stat | DF | p-Value | CI Lower | CI Upper |
| --- | --- | --- | --- | --- | --- | --- | --- |
| (Intercept) | 1.291400 | 0.44894 | 2.8764 | 115 | 0.00479 | 0.40208 | 2.1806 |
| WMC | 0.083428 | 0.05040 | 1.6553 | 115 | 0.10058 | -0.01640 | 0.18326 |
| Inattentive | -0.034875 | 0.01630 | -2.1396 | 115 | 0.03450 | -0.06716 | -0.00259 |
| Hyperactive | 0.031835 | 0.02144 | 1.485 | 115 | 0.14027 | -0.01063 | 0.07430 |

Random effects covariance parameters:

| Group | Name1 | Name2 | Type | Estimate |
| --- | --- | --- | --- | --- |
| Exp (4 Levels) | (Intercept) | (Intercept) | std | 0.23505 |

| Group | Name | Estimate |
| --- | --- | --- |
| Error | sqrt(Dispersion) | 0.11232 |

